# A scalable approach to inoculate plant viral vectors into plant tissue using non-pathogenic, transgenic galls

**DOI:** 10.64898/2026.02.16.706162

**Authors:** Stacy L. DeBlasio, Feng Gao, Zhiqian Pang, David O. Igwe, Samantha R. Sullivan, Yu-Hui Wang, Marco Pitino, Samuel Coradetti, Anne Simon, Michelle Heck

## Abstract

Vascular plant pathogens transmitted by insects inflict devastating economic losses on crops worldwide. By obstructing and usurping the natural flow of nutrients, plant infection by these pathogens drastically reduces yields, vigor and productivity. Treating fruit-bearing trees against persistent vascular pathogens poses a unique challenge, as systemic delivery of therapeutics must navigate the compartmentalized architecture of the tree’s vascular system under changing environmental conditions without disrupting fruit production or long-term tree health. Plant viruses have gained traction as a novel approach to deliver therapies to crop plants, including fruit trees, but delivery of viral vectors to orchards at scale remains a significant challenge. We tested whether transgenic galls can be used to systemically infect plants with a plant virus infectious clone. We combined the plant growth regulator gene cassette from *Agrobacterium tumefaciens* strain C58 with the wild-type strain of citrus yellow vein associated virus-1 (CY1) on a single plasmid within the T-DNA for plant cell transformation. Using EHA105, a disarmed strain of *A. tumefaciens*, we inoculated stems with these gall-forming plasmids and initiated systemic CY1 infection in citrus and *Arabidopsis thaliana* over three independent experiments. We provide proof-of-concept that transgenic galls, referred to as symbionts, can launch the systemic infection of CY1 in economically important and model plants. Symbiont delivery of therapeutic viral vectors is theoretically scalable from inoculation of mother trees within the nursery to millions of trees in the field and may be a valuable tool for the commercial delivery of therapeutic plant viral vectors.

## Text

Pathogens that infect the xylem and/or phloem network (i.e., vasculature) of crop trees, causing widespread damage to these plants by compromising water and nutrient transport. This disruption leads to a weakened root system, canopy dieback, decreased fruit set/ripening and in severe cases, death of the tree. Several vascular plant diseases are currently causing significant economic losses in the United States (US) and globally. Notable examples include Cherry X-disease and pear decline, caused by the phytoplasma *‘Candidatus Phytoplasma pruni’* (Wright et al., 2021) and *Ca. Phytoplasma pyri*, respectively (Cui et al., 2025); Panama disease of bananas, caused by the fungus *Fusarium oxysporum* (Aguilar-Hawod et al., 2019); Pierce’s disease of grapevines, caused by the xylem-limited bacterium *Xylella fastidiosa* (Davis et al., 1978); and citrus greening disease (Huanglongbing, HLB), regarded as the most severe threat to citrus production worldwide (Zheng et al., 2024).

HLB is associated with plant infection by a fastidious, gram-negative bacterium in the genus ‘*Candidatus Liberibacter’*. In parts of the world where citrus is grown, including the US, Asia and South America, the predominant and most virulent species is ‘*Candidatus* Liberibacter asiaticus’ (*C*Las) (Younas et al., 2025). Transmitted directly to citrus phloem by its sap-sucking insect psyllid vectors in the Hemiptera, the *C*Las bacterium has been shown to induce a hyperactive plant immune response that produces high levels of reactive oxygen species (ROS) and increased callose/P-protein accumulation that leads to severe oxidative stress and blockage of nutrient flow in the phloem, respectively (Ma et al., 2022). Observable symptoms of the disease start with asymmetric, yellow mottling on leaves (a key diagnostic symptom), vein corking, and stunted root growth, which is followed by more severe declines in vigor such as twig dieback, root rot and leaf defoliation (Tipu et al., 2021). Symptoms in fruit include lopsided growth, smaller than normal size, uneven ripening where bottoms remain green, and bitter juice (Dala-Paula et al., 2018). Fruit drop prior to harvest, where the calyx abscission zone or “button” that connects the fruit to the tree becomes weakened, is also a common symptom of *C*Las-infected citrus that compounds economic losses in Florida during severe weather events (Chen et al., 2016).

Although progress has been made to sustain citrus tree productivity under HLB pressure, including HLB-tolerant rootstock and scions as well as the use of therapeutic trunk injection (Archer et al., 2023; Heck et al., 2025), research efforts have failed to solve the problem economically. Antimicrobial peptides (AMPs) and RNA interference signaling molecules can disrupt *C*Las infection *in planta* (Higgins et al., 2024; Huang et al., 2021; Killiny et al., 2025; Padilla et al., 2025), *C*Las acquisition by psyllids (Higgins et al., 2024; Huang et al., 2021), and induce psyllid mortality (El-Shesheny et al., 2013; Liu et al., 2020). However, next generation biological-based pesticides are costly to produce at scale. Furthermore, *C*Las is a phloem-restricted pathogen and delivery of peptides and RNA therapies to the phloem requires delivery technologies that target plant vascular tissues (Archer et al., 2023; Ojo et al., 2024). Although devices used for trunk injections have been developed to target the vasculature (xylem and phloem) (Ojo et al., 2024; Rhodes et al., 2023) and are commercially available to growers (Heck et al., 2025), the process still relies on wounding the tree, which the tree immediately repairs eventually blocking further treatment uptake. Several years of data in Florida demonstrate the safety of trunk injection over a multi-year time scale (Archer et al., 2023; Heck et al., 2025), but the long-term impacts of trunk injection on tree health remain unknown.

The use of “disarmed” (i.e., symptomless) plant viruses and viral replicons to produce peptides, proteins, or siRNAs that inhibit pathogens, enhance crop traits such as drought resistance, or target insect vectors has gained traction and represent a biologically-based strategy for delivering therapies to plants (Hajeri et al., 2014; Shen et al., 2024; Torti et al., 2021; Venkataraman & Hefferon, 2021), including HLB treatments to citrus trees. For example, a mild strain of the phloem-limited, *citrus tristeza virus* (CTV) has been developed as a peptide expression vector and gene silencing tool in citrus and psyllids (Hajeri et al., 2014; Ibanez et al., 2023; Killiny et al., 2025). Additionally, the small, phloem-limited, umbravirus-like virus, citrus yellow vein associated virus 1 (CY1, also known as CYVaV) (Kwon et al., 2021; Ying et al., 2024), is an advantageous alternative viral delivery vector for citrus therapeutics. The tiny 2.7 kb CY1 genome codes for viral replication proteins and lacks a capsid protein, canonical movement protein and silencing suppressor proteins that are generally required for other RNA plant viruses to successfully infect their host and to be transmitted (Kwon et al., 2021). In the absence of a helper virus from the luteo-, enamo-, or polerovirus genera, CY1 relies on the phloem-mobile plant protein PP2 to move systemically within plants (Ying et al., 2024). CY1 can be modified into a stable, effective virus-induced gene silencing (VIGS) vector in the *Nicotiana benthamiana* and transferred to Mexican lime using dodder, a parasitic plant or by vacuum infiltration into leaves with *Agrobacterium tumefaciens* (Ying et al., 2024). However, no scalable approach currently exists to inoculate thousands or even millions of citrus trees with CTV, CY1, or any other therapeutic plant viral vectors.

Inspired by the molecular biology of crown gall formation on plants, we developed a method to engineer non-pathogenic galls produced by *A. tumefaciens* to express recombinant genes on-planta (Heck et al. 2026, Shatters et al., 2021). These galls are living structures that consist of autonomously dividing plant cells vascularly connected to the main plant host (Aloni et al., 1995; Pradel et al., 1999; Ullrich & Aloni, 2000). The development of crown galls, caused by *A. tumefaciens* and other bacterial species, is a well-studied process that involves expression of plant growth regulator (PGR) genes found within the T-DNA region of the bacterium’s Ti plasmid following plant cell transfection (Akiyoshi et al., 1984; Inze et al., 1984; Kober et al., 1991; Schroder et al., 1984). Stable incorporation of the plant growth regulator genes into the genome of a subset of plant cells triggers the formation of an “organ-like” growth that protrudes from the host stem, resulting in the gall. In pathogenic crown gall infections, genes encoding the production of opines are also transferred into the plant genome within the T-DNA region (Koncz et al., 1983). Opines are a class of carbohydrate derivatives used by *Agrobacterium* as a carbon source (Kim et al., 2008). Other genes regulating the catabolism and uptake of specific opines are encoded on the Ti plasmid (Gordon & Christie, 2014). Thus, the T-DNA and Ti plasmid encode genes that have been favored by selection to generate a specialized niche for *Agrobacterium* infection and survival.

By developing a plant transformation vector with a T-DNA that a) harbors the PGR cassette, b) is devoid of the opine-producing genes and c) expresses a gene of interest (in this case the CY1 virus), we developed a molecular approach to turn pathogenic galls into functional, on-planta, bio-factories, which we refer to a “symbiont” to reflect its non-pathogenic nature and potential beneficiary effects on the host plant (Heck et al., 2026; Shatters et al., 2022). A currently pending patent application (Shatters et al., 2022) further defines the concept more technically, stating that a “symbiont” is a plant cell or plurality of plant cells comprising a polynucleotide encoding a phytohormone biosynthetic enzyme (e.g., at least one polynucleotide encoding one or more phytohormone biosynthetic enzymes and a polynucleotide of interest). Vectors with the PGR cassette and gene of interest are referred to as pSYM (Fig. 1). The pSYM vectors contain a four-gene PGR cassette from *A. tumefaciens* strain C58 (*I3L, IaaH, IaaM, Ipt*) driven by the native C58 promoter(s) for this gene cluster (Akiyoshi et al., 1983; Joos et al., 1983; Otten et al., 1999). The products of these genes induce “symbiont”-type gall formation from tissues in the stem by establishing auxin and cytokinin gradients that regulate cell division and vascular tissue differentiation. The full sequence of pSYM is available on GeneBank (PQ563460; Heck et al. 2026).

**Figure 1:**
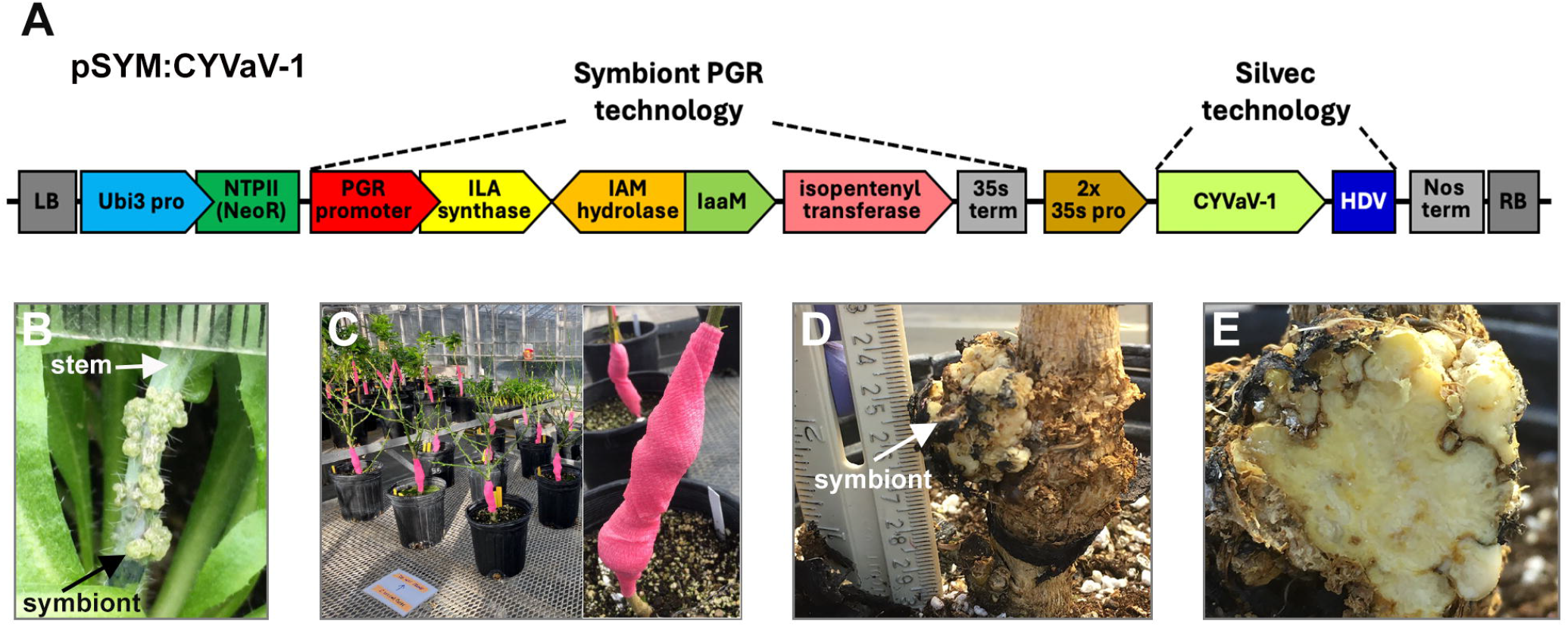
The pSYM vector produces gall-like structures on citrus fruit-bearing tree stems that naturally integrate with the vascular system of a host plant and expresses CY1. **(A)** Schematic of the symbiont-forming PGR cassette and viral expression region of the pSYM:CY1 plasmid used in this study. The construct was made by cloning a 4-gene PGR cassette: *I3L* (ILA synthase), *IaaH* (IAM hydrolase), IaaM (tryptophane 2-monooxygenase), and *Ipt* (isopentyltransferase) from *A. tumefaciens* strain C58 with its endogenous promoter (PGR promoter) into the plant binary vector pUSHRL-26 (Krystel et al., 2021) upstream of the double 35S CaMV promoter. Downstream of the 2x 35S promoter, the sequence encoding CY1 with a hepatitis delta virus (HDV) ribozyme sequence was added for constitutive expression of CY1 within symbionts and enhanced delivery to the phloem of plants. LB = left T-DNA border sequence; Ubi3 pro = ubiquitin 3 promoter; NTPII = the first exon and intron of ubiquitin 3 fused to the plant Kan resistance gene NeoR; 35S term = 35S terminator from CaMV; Nos term = Nos gene terminator’ RB = right T-DNA border sequence. (**B**) Photo shows characteristic symbiont growth on an *Arabidopsis* inflorescence stem at 25+ days post infiltration with pSYM. Ruler indicates millimeters. (**C**) Images show outer pink vet tach wrapping of citrus symbionts growing on the stem of defoliated 1 to 2-year greenhouse grown citron trees at one month post-symbiont inoculation. Inset shows closer view of symbiont protrusion under wrapping at ∼4 to 5 months post-symbiont inoculation **(D)** Side view of an unwrapped pSYM:CY1 expressing citrus symbiont at ∼ 7 months post-symbiont inoculation. Ruler indicates centimeters. **(E)** Inside tissue of same symbiont with distinct niches of dividing cells (clear and yellowed) and developed vasculature (white).

To test whether symbionts induced on stems via pSYM can be used to systemically infect plants with plant viral vectors, we engineered the pSYM vector to express the infectious clone of CY1. We cloned the CY1 genome followed by a HDV ribozyme sequence (Ying et al., 2024) into the T-DNA region of our symbiont-inducing pSYM vector downstream of a double 35S promoter from cauliflower mosaic virus (CaMV) for constitutive expression in symbiont cells (Fig. 1A). The full sequence of the pSYM vector expressing CY1, referred to as pSYM:CY1, is available on GenBank (PX738482). The pSYM vector also contains kanamycin resistance genes for selection in bacteria (KanR in the vector backbone) and in transformed plant cells (NeoR in the T-DNA). The complete, sequence-verified pSYM:CY1 vector was transformed into the disarmed *A. tumefaciens* lab strain EHA105 using electroporation and positive colonies selected for on kanamycin supplemented (100 μg/mL) Lennox Broth agar plates at 28 °C. Four independent, kanamycin-resistant pSYM:CY1 EHA105 colonies were chosen and assayed for symbiont formation and for the ability to induce CY1 systemic infection. We used the model plant *Arabidopsis thaliana* (ecotype: Columbia) and greenhouse grown citron trees for the experiments.

*A. thaliana* plants support robust symbiont formation following stem inoculation with pSYM vectors (Fig. 1B). Thus, we inoculated 3.5 to 4-week-old *A. thaliana* (ecotype: Columbia) with pSYM:CY1, pSYM:GFP (CY1 negative control), pCY1 (the non-symbiont forming pCB301-CY1 vector) with or without the viral silencing suppressor P14. *Agrobacterium* strains were grown overnight at 28 °C with shaking (200 rpms). Bacterium was pelleted by a 25 min centrifugation (up to 10,000 RCF) at 24 to 26 °C and washed two times in infiltration buffer (10 mM MES, pH 5.6; 10 mM MgCl_2_; 400 μM acetosyringone). Two *Agrobacterium* isolates of pSYM:CY1 were tested and six plants were inoculated per construct (there was some loss of biological replicates due to random inflorescence breakage during the growing process). Final inoculation culture ODs were between 1.0 and 1.1 for each strain in fresh buffer, even when pCY1 and P14 were mixed 1:1 for co-inoculation. Inflorescence stem lengths were between 1 and 8 mm and inoculated at three points longitudinally with droplets of culture at the end of an insulin syringe. Plants were grown in the Heck Lab greenhouse in Ithaca, New York on the campus of Cornell University during winter and maintained at temperatures ranging from 70 to 75 °F. At 49 days post-inoculation, symbiont growth was evident in all pSYM-inoculated plants but absent in pCY1-inoculated plants (data not shown). To assess systemic movement of CY1, tissue was collected from symbionts or inoculation sites (pCY1 plants) and from distal tissues including stem above symbionts, roots, cauline leaves, and green siliques. Total RNA was extracted (Synergy™ 2.0 kit with on-column DNase I treatment), and cDNA synthesized from 500 ng of RNA using the Superscript III First-Strand Synthesis System using random hexamers. cDNA was diluted 1:5 and used for PCR detection of CY1 vRNA with primers CY-1634-1652-F (5’-GAAGGCGTCACGTCCGTGT-3’) and CY-2470-89-R (5’-CCACAGTGCTATCGCTCCAA-3’), producing an 856 bp amplicon. We performed PCR with Phusion™ polymerase (20 µL reactions, Tm = 55 °C, 30 s extension).

Inoculation of *A. thaliana* with pSYM:CY1 resulted in systemic infection of *A. thaliana* with CY1 (note that CY1 has a wide host range likely due to its use of a highly conserved host movement protein). Of eight plants infiltrated with pSYM:CY1, six contained CY1 vRNA expression in symbiont tissue (75%) (Fig. 2A). In five plants inoculated with pSYM:CY1 (62.5%), the virus was detected in stems, roots, or both. In contrast, among seven plants infiltrated with the non-symbiont pCY1 clone (with or without silencing suppressor P14), no virus was detected in any tissue of four plants including the inoculation site, and the virus was sporadically detected in the areal tissues of three plants, with one detection limited to the inoculation site. Co-inoculation with P14 had no effect on virus detection. Notably, pCY1 plants did not show CY1 presence in roots, whereas three pSYM:CY1 plants did. No amplification of the CY1 856 bp amplicon was observed in pSYM:GFP controls, confirming primer specificity.

**Figure 2:**
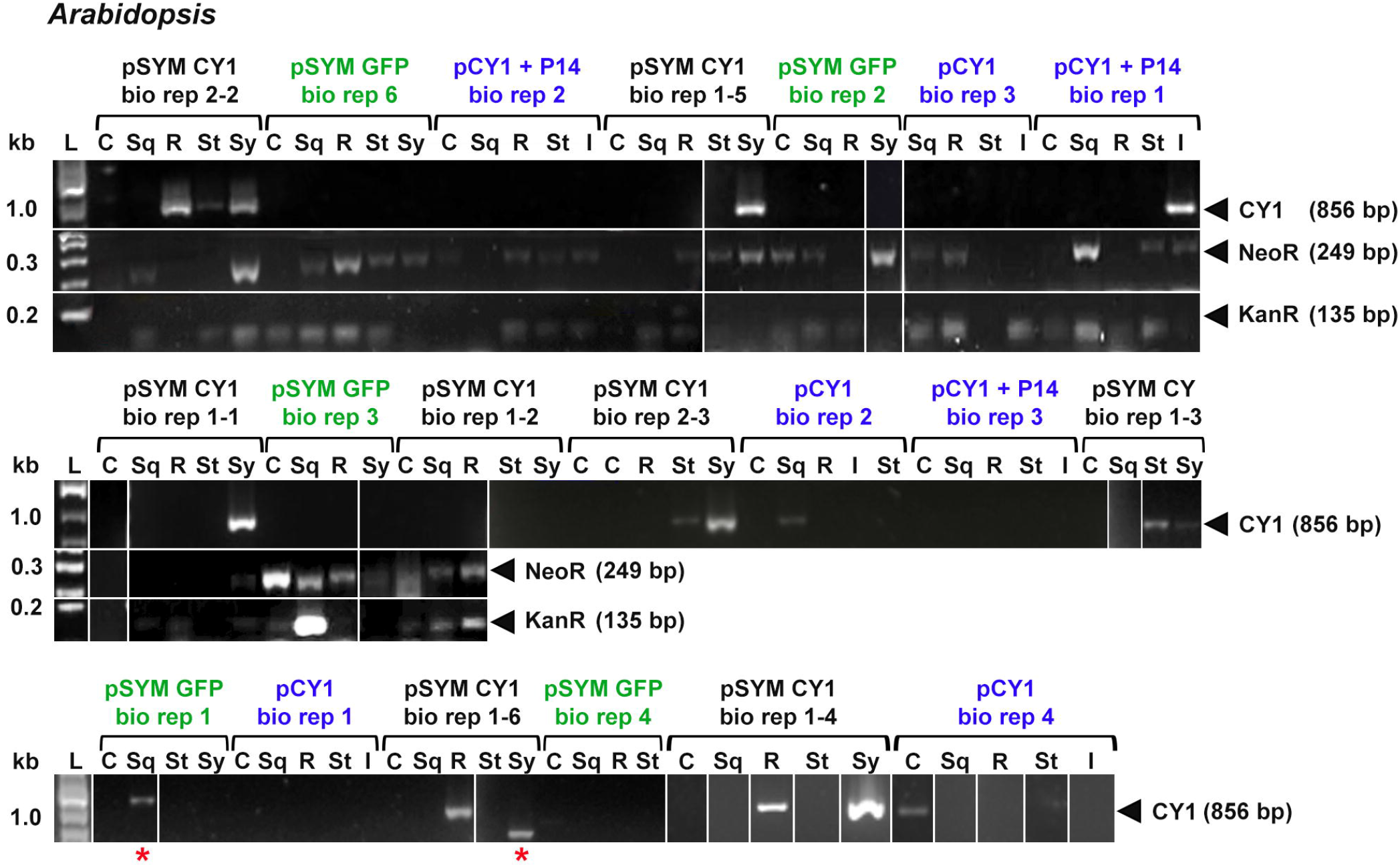
Efficient systemic delivery of CY1 to model plant species *Arabidopsis thaliana* using the pSYM vector is dependent on symbiont development. RT-PCR detection of CY1 vRNA amplicon (856 bp) in systemic tissues of Arabidopsis 49 days post-infiltration with pSYM:CY1 (isolate 1 or 2, black), pSYM:GFP (green) or the non-symbiont forming infectious clone pCB301-CY1 (pCY1), without or with silencing suppressor P14 (blue) using *A. tumefaciens* strain EHA105. Detection of amplicons corresponding to the bacterial and plant-expressed kanamycin resistance genes, KanR (135 bp) and NeoR (249 bp), respectively, was also assessed for some of the biological replicate plants. L = GeneRuler 1 kb Plus DNA ladder (Thermo Fisher Sci); C = cauline leaves; Sq = immature siliques: R = roots; St = stem 2+ mm above symbiont/inoculation site; Sy = symbiont tissue; I = inoculation site stem tissue. Tissue samples that are missing were lost due to technical reasons. PCR reactions were run on three separate gels with all controls and ladders. Final image in panel E was created by manually grouping together the results. Wells that were not sequential are indicated by white lines.

Systemic movement of *A. tumefaciens* has been reported in some plants (Cubero et al., 2006). While the lack of systemic infection of CY1 in plants inoculated with pCY1 supports the hypothesis that systemic movement of CY1 vector-harboring *A. tumefaciens* cells from the inoculation site was not responsible for the systemic detection of CY1, we wanted to rule out the possibility by assaying for *A. tumefaciens* genes in systemic plant tissues. We used RT-PCR for the detection of kanamycin resistance genes from the pSYM and pCB301 plasmids using bacterial KanR-F2/R2 and plant NeoR-F1/R1 primer sets (KanR-F2: 5’-AGACGGAAAAGCCCGAAAGAG-3’ and KanR-R2: 5’-TGTCATACCACTTGTCCGCC-3’; plant NeoR-F1: 5’-CAAGATGGATTGCACGCAGG-3’ and plant NeoR-R1: 5’-TTCAGTGACAACGTCGAGC-3’), to monitor potential *A. tumefaciens* movement/persistence, mRNA expression, or T-DNA integration in distal tissues (Fig. 2). KanR and NeoR cDNA amplicons from pSYM and pCB301 were detected discordantly across tissues even in pSYM:GFP controls, suggesting either persistence of “disarmed” *Agrobacterium* and/or movement of vector-associated mRNA but without congruent CY1 detection. While our results cannot completely rule out *A. tumefaciens* movement in *Arabidopsis*, CY1 was consistently detected along a tissue gradient when plants were infiltrated with pSYM:CY1 but not when inoculated with pCY1. These results support the hypothesis that symbiont formation enhances systemic delivery of CY1 in *A. thaliana*, especially to roots.

We also tested CY1 delivery by symbionts on citrus. As mentioned above, CY1 infection from the non-symbiont forming pCY1 infectious clone has only been demonstrated in Mexican lime using dodder, vacuum infiltration of *A. tumefaciens* into leaves, or traditional graft inoculation which takes 12-15 months for systemic CY1 to be detected (Ying et al., 2024). Our symbiont inoculation protocol for citrus is similar to the one used for *Arabidopsis*, with the following exceptions: 20 μL of an OD_600_ ≅ 1.0 infiltration culture of EHA105 harboring pSYM:CY1 was added to four biopsy cut-outs (4 mm) located in a spiral pattern along the main stem of 1 to 2-year-old healthy or *C*Las-infected, greenhouse grown, citron trees. A set of *C*Las-infected trees were also inoculated with 20 μL of EHA105 harboring the non-symbiont forming pCY1 vector with or without P14. Inoculation sites were immediately wrapped in two layers of parafilm, followed by one layer of black Frost King flex foam, and one layer of light color vet tack, to keep humidity levels high around the inoculation site, which is a key requirement for sustained symbiont growth on citrus (Fig. 1C). Representative images of one of these symbionts on citron is shown in Fig. 1D and E.

On healthy citron, we tested four EHA105 clones harboring pSYM:CY1 (denoted pSYM:CY1 isolate 1-4), using three trees inoculated per isolate for a total of 12 trees. We used a GFP-expressing symbiont vector a negative control. At one-month post-symbiont inoculation, plants were completely defoliated and the canopy allowed to re-grow to potentially provide a stronger sink than the symbiont. At four months post-symbiont inoculation, leaf discs from three canopy positions were assayed for CY1 vRNA by RT-PCR following the same protocol outlined above (Fig. 3A). We detected CY1 starting at the bottom of the canopy, in at least one biological replicate per pSYM:CY1 isolate, which was four out of 10 total trees (40%). By six months post-symbiont inoculation, 90% of pSYM:CY1 trees showed CY1 vRNA in at least one canopy site and roots, with 60% trees positive at multiple canopy positions (Fig. 3B), indicating progressive systemic spread from four to six months post-symbiont inoculation. Two trees inoculated with isolate 3 died before four months post-symbiont inoculation, likely due to small stem diameters (<10 mm) which we have observed to occasionally be insufficient to support the combination of potted trees (data wounding with a 4 mm biopsy punch and rapid symbiont growth in greenhouse not shown).

**Figure 3:**
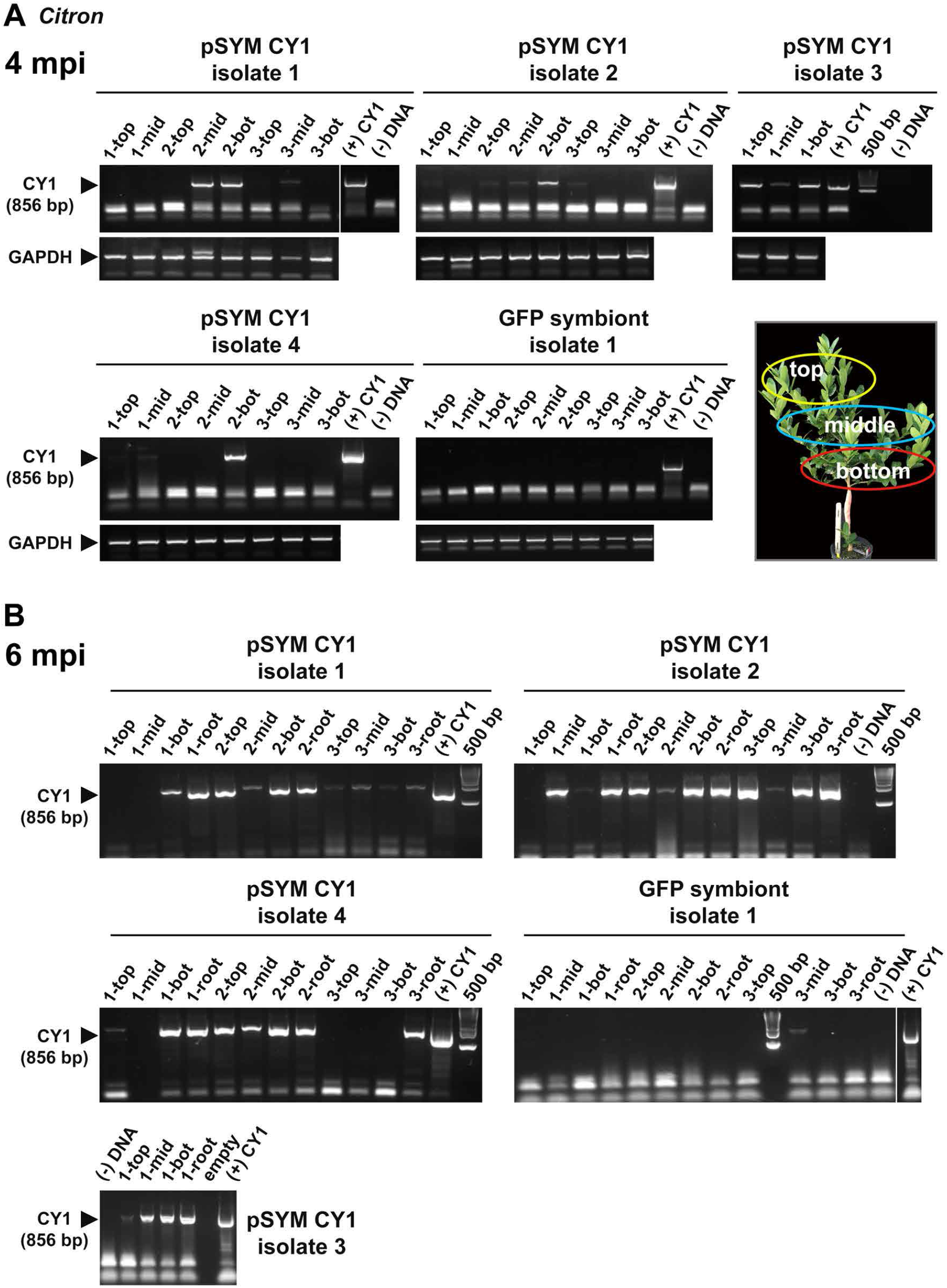
Successful systemic delivery of CY1 via symbiont technology to the leaf canopy and roots of healthy citron trees. RT-PCR detection of the 856 bp amplicon specific to the CY1 genome in canopy leaves or roots of greenhouse grown, *C*Las negative (i.e., healthy) citron trees at **(A)** 4 and **(B)** 6 months post-stem inoculation with four different *Agrobacteruim* clone isolates of pSYM:CY1 or pSYM:GFP. Amplification of the endogenous control (GAPDH gene) was also assessed as a technical control in these same samples. Image shows canopy position of leaf pools (3 leaves pooled per sample) from which total RNA was extracted: top of canopy (top, yellow circle); middle of canopy (mid, blue circle) and bottom of canopy (bot, red circle). The bottom of the canopy is where the most mature leaves (oldest) reside on a branch. Wells are labeled as biological replicate number-tissue position. (+) CY1 = infectious clone plasmid as positive PCR control; (-) DNA is negative PCR control without any cDNA added; 500 bp DNA ladder (brightest band = 500 bp). Black arrowhead indicates specific amplicon position.

To assay for feasibility of pSYM to deliver plant viral vectors as *C*Las-targeted therapies, we tested whether pSYM:CY1 can systemically infect *C*Las-infected citron. We inoculated three, 1 to 2-year-old, *C*Las positive (Ct ≤ 33), citron trees with pSYM:CY1-1, pCY1 ± P14 or pSYM:GFP following the same protocol used for healthy citron. At seven months post-symbiont inoculation, 4 mm leaf discs from three mature leaves (two discs per leaf) were pooled from each of four different canopy quadrants (top, right, left and bottom) and analyzed for the presence of CY1 vRNA using RT-PCR following total nucleic (TNA) extraction protocol used in Higgins et al. (2024) with a subsequent DNase treatment and cDNA synthesis following the *Arabidopsis* protocol above. All plants inoculated with pSYM:CY1 had at least two canopy quadrants test positive for CY1 (Fig. 4A). One biological replicate, pSYM:CY1-2, tested positive in all four quadrants. Out of the six trees inoculated with the non-symbiont forming pCY1 clone, one quadrant of one plant initially tested positive for CY1. However, extensive re-sampling of this tree resulted in no further detection of CY1 indicating the initial result was a false-positive due to technical contamination. Therefore, our data shows that CY1 systemic movement in citron could only be established from constitutive expression from the symbiont and not just by application/uptake of *Agrobacteria* harboring pCY1 through a wound in the stem. *C*Las titer in the leaf canopy on average was slightly lower than pSYM:GFP plants (Fig. 4B), but the titer difference was not statistically significant (Student T-test, *p*-value = 0.21). To assess whether CY1 affected symbiont growth, we compared pSYM:CY1 and pSYM:GFP symbionts and found no differences in symbiont size or development frequency (data not shown). Thus, CY1 infection does not adversely affect symbiont development on citrus, an important finding supporting the potential use of symbionts for CY1 delivery in future commercial applications. Visual symptoms of CY1 infection in citron were not detected on the CY1-infected plants in the timeframe of our experiment (data not shown).

**Figure 4:**
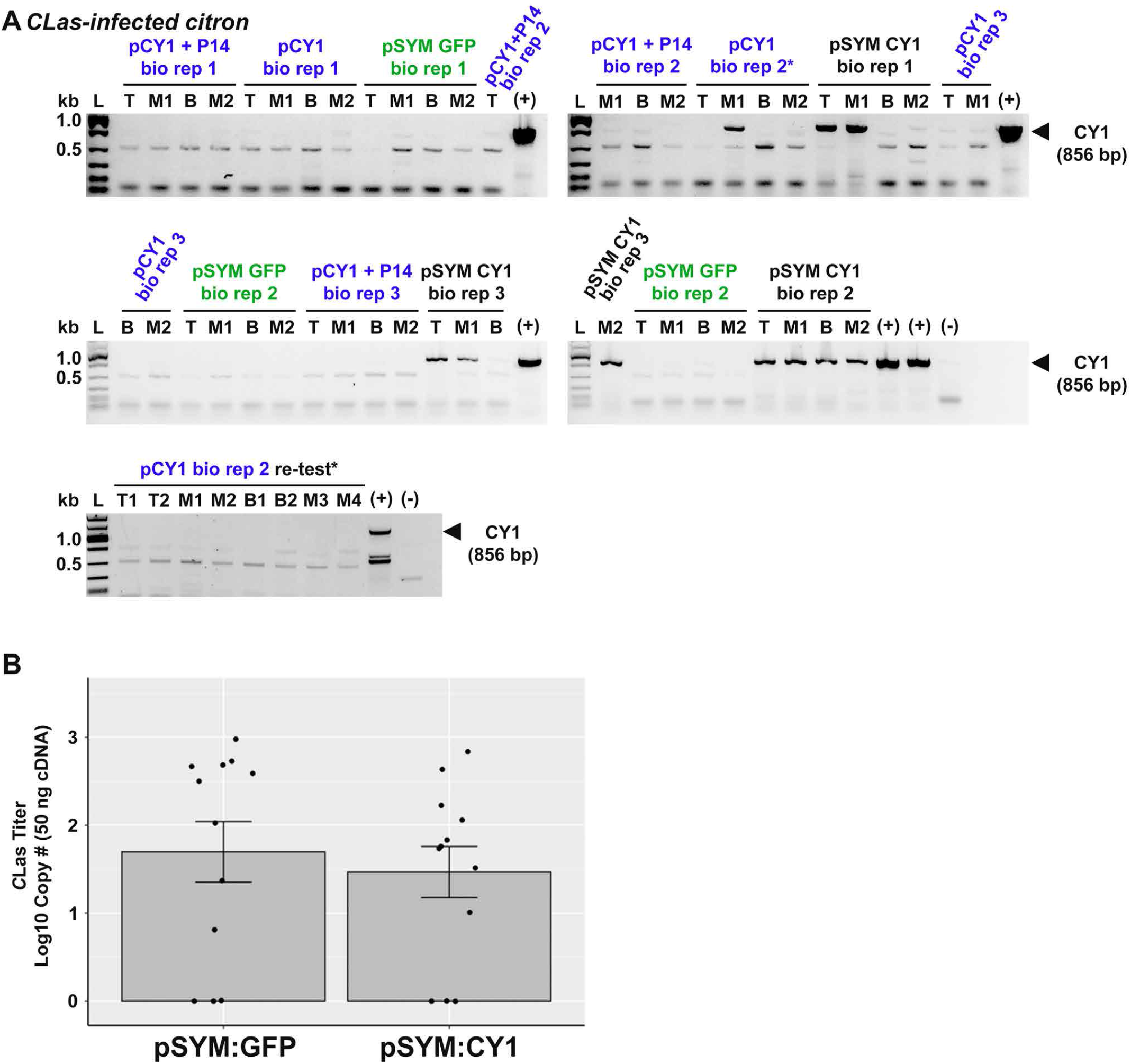
Systemic delivery of CY1 via symbiont technology to the leaf canopy of *C*Las-infected citron trees. **(A)** RT-PCR detection of the 856 bp amplicon specific to the CY1 genome in canopy leaves of greenhouse grown, *C*Las-positive citron trees at ∼7 months post-stem inoculation with pSYM:CY1 (isolate 1, black), pSYM:GFP (green) or the non-symbiont forming infectious clone pCB301-CY1 (pCY1) with or without P14 (blue). Total RNA was extracted from pools of three leaves per canopy quadrant position: T = top of canopy; M1 = middle, right; B = bottom of the canopy; and M2 = middle, left. L = Fast DNA Ladder (New England Bio). Blue star indicates biological replicate tree infiltrated with non-symbiont forming pCY1 that had one leaf quadrant test positive for CYVaV-1. Black starred gel shows RT-PCR re-analysis of eight additional leaf pools from that same plant, all of which tested negative indicting first result was due to technical contamination. **(B)** Bar graph shows quantitative qPCR analysis of *C*Las titer (Higgins et al., 2024) in same leaf samples of trees inoculated with pSYM:CY1 or pSYM analyzed in panel A. Titer is given as Log10 copy number detected in 50 ng of cDNA relative to a qPCR standard dilution of a plasmid clone of the *C*Las 16s RNA amplicon. Black dots represent a leaf pool sample per canopy quadrant, n = three biological replicate trees per construct. Error bars represent standard error.

In conclusion, we demonstrate across three independent experiments, that CY1, a phloem-restricted umbra-like virus, can be expressed in symbiont tissue and delivered to systemic tissues in *Arabidopsis* and citron using the gall-forming pSYM vector. Notably, systemic plant infection from the symbiont tissue occurred in the absence of a helper virus, coat protein, classical movement protein, or viral silencing suppressor. In greenhouse grown citron, systemic CY1 infection had no effect on *C*Las titer or HLB symptom development. We showed that effective delivery of the virus is dependent on symbiont growth, as direct application of a non-symbiont forming infectious clone of CY1 to plant stems did not result in sustained systemic infection, especially to roots. Detection of Kan mRNA from the vectors in *Arabidopsis* tissues did not directly coincide with CY1 expression in those same samples (Fig. 2A), suggesting limited or no contribution from any possible *Agrobacterium* systemic dissemination to viral systemic spread. Our work lays the foundation for symbionts to be used as a scalable delivery strategy for CY1 vectors in citrus, which has already shown to be an excellent, stable, virus-induced gene silencing (VIGS) vector in *N. benthamiana*. Additional research is ongoing to determine whether symbionts can be used to inoculate plants with larger plant viruses, such as CTV. Virally delivered siRNAs or peptides targeting plant immunity genes or *C*Las, respectively, from symbionts, may be used as a feasible strategy to protect nursery trees from future infection or to lower bacterial titer, mitigate disease symptoms or boost tree health when applied to *C*Las-infected trees in the field. Symbionts may be useful to deliver viral vectors to a range of perennial plant species.

## Acknowledgements

This research was funded by the USDA National Institute of Food and Agriculture Emergency Citrus Disease Research and Extension Program award number: 2022-70029-38503 and the USDA Agricultural Research Service project numbers: 8062-22410-0007-000D and 8062-30400-001-00D.

## Bibliography

Aguilar-Hawod, K. G. I., de la Cueva, F.M., & Cumagun, C. J. R. 2019. Genetic diversity of Fusarium oxysporum f. Sp. Cubense causing panama wilt of banana in the philippines. Pathogens. 9: 32.

Akiyoshi, D. E., Klee, H., Amasino, R. M., Nester, E. W., & Gordon, M. P. 1984. T-DNA of Agrobacterium tumefaciens encodes an enzyme of cytokinin biosynthesis. Proc. Natl. Acad. Sci. U.S.A. 81: 5994–5998.

Akiyoshi, D. E., Morris, R. O., Hinz, R. S. M. B.; Kosuge, T., Garfinkle, D. J., Gordon, M. P., & Nester, E. W. 1983. Cytokinin/auxin balance in crown gall tumors is regulated by specific loci in the T-DNA. Proc. Natl. Acad. Sci. U.S.A. 80: 407–411.

Aloni, R., Pradel, K. S., & Ullrich, C. L. 1995. The three-dimensional structure of vascular tissues in Agrobacterium tumefaciens-induced crown galls and in the host stems of Ricinus communis l. Planta. 196: 597–605.

Archer, L., Kunwar, S., Alferez, F., Batuman, O., & Albrecht, U. 2023. Trunk injection of oxytetracycline for Huanglongbing management in mature grapefruit and sweet orange trees. Phytopathology. 113: 1010–1021.

Chen, H., McCollum, G., Baldwin, E., & Bai, J. 2016. Impacts of Huanglongbing symptom severity on fruit detachment force and mechanical properties of sweet oranges (Citrus sinensis). HortScience. 51: 356–361.

Cubero, J., Lastra, B., Salcedo, C. I., Piquer, J., & Lopez, M. M. 2006. Systemic movement of Agrobacterium tumefaciens in several plant species. J. Appl. Microbiol. 101: 412–421.

Cui, W., Alburquenque, C., Pacini, F., Gonzalez, C., Bianco, P., Cabrera, S., Llanten, T., Fuentes, J., Gamboa, C., Bertaccini, A., Fiore, N., & Zamorano, A. 2025. Draft genome of ‘Candidatus Phytoplasma pyri’ and phylogenetic diversity among chilean and italian strains. Phytopathology. 115: 1080–1085.

Dala-Paula, B. M., Plotto, A., Bai, J., Manthey, J. A., Baldwin, E. A., Ferrarezi, R. S., & Gloria, M. B. A. 2018. Effect of Huanglongbing or greening disease on orange juice quality, a review. Front .Plant. Sci. 9: 1976.

Davis, M. J., Purcell, A. H., & Thomson, S. V. 1978. Pierce’s disease of grapevines: Isolation of the causal bacterium. Science. 199: 75–77.

El-Shesheny, I., Hajeri, S., El-Hawary, I., Gowda, S., & Killiny, N. 2013. Silencing abnormal wing disc gene of the asian citrus psyllid, Diaphorina citri disrupts adult wing development and increases nymph mortality. PLoS One. 8: e65392.

Gordon, J. E., & Christie, P. J. 2014. The Agrobacterium Ti plasmids. Microbiol. Spectr. 2:

Hajeri, S., Killiny, N., El-Mohtar, C., Dawson, W. O., & Gowda, S. 2014. Citrus tristeza virus-based RNAi in citrus plants induces gene silencing in Diaphorina citri, a phloem-sap sucking insect vector of citrus greening disease (Huanglongbing). J. Biotechnol. 176: 42–49.

Heck, M. L., Larson, N. R., Locatelli, G., Cochrane, E. F., Coradetti, S., Hallman, L., Johnson, L., Estes, W. C., Makar, A., Demirden, N., Pitino, M., Shatters, R. G., Adair, R. C., Giles, F., Fox, J.-P., Stuehler, D., Hodge, J., Hoffman, J., Blake, V., Ulysse, L., Ramirez-Barrera, L., McKenna, R., Thompson, L., Bennett, L., Larrea-Sarmiento, A., Olmedo-Velarde, A., Barkee, S., Weeks-Purdy, C., Zambon, F., Shende, K., Rossi, L., D’Elia, T., Ritenour, M. A., Scully, B. T., & Niedz, R. P. 2025. The “Grove-First” framework: Starting in the grove to find therapies for Huanglongbing. Plant Disease. doi: 10.1094.PDIS-07-25-1379-RE.

Heck, M.L., Pitino, M., Coradetti, S., DeBlasio, S.L., Cooper, W.R., Shrum, L., Harper, D., Stallone, M., Aspen, S., Cook, R., Rhodes, B., Sullivan, S., Schechter, E., Cochrane, E., Larson, N., Locatelli, G., Hodge, J., Grando, M.F., Wang, L., Ariyarante, M., Tibebu, Strange, R., Howe, K.J., Makar, A., Stuehler, D., Thompson, L., Shende, K., Hentz, M., Gaza, N., Weeks-Purdy, C., Chang, B., Nikoomanzar, A., Bennett, L., Demirdenn, S., Hunter, W.B., Thomson, J., Ritenour, M., Rossi, L., Cano, L.M., Adair, R.C. Jr., Stover, E., McKenzie, C., Niedz, R. & Shatters, R.G. Jr. 2026. A new use of Agrobacterium plant growth regulator genes for plant bioengineering. Frontiers in Plant Science. In press.

Higgins, S. A., Igwe, D. O., Coradetti, S., Ramsey, J. S., DeBlasio, S. L., Pitino, M., Shatters, R. G., Jr., Niedz, R., Fleites, L. A., & Heck, M. 2024. Plant-derived, nodule-specific cysteine-rich peptides as a novel source of biopesticides for controlling citrus greening disease. Phytopathology. 114: 971–981.

Huang, C. Y., Araujo, K., Sanchez, J. N., Kund, G., Trumble, J., Roper, C., Godfrey, K. E., & Jin, H. 2021. A stable antimicrobial peptide with dual functions of treating and preventing citrus Huanglongbing. Proc. Natl. Acad. Sci. U.S.A. 118: e2019628118

Ibanez, F., Vieira Rocha, S., Dawson, W. O., El-Mohtar, C., Robertson, C., Stelinski, L. L., & Soares-Costa, A. 2023. Gene silencing of cathepsins B and L using CTV-based, plant-mediated RNAii interferes with ovarial development in Asian citrus psyllid (ACP), Diaphorina citri. Front. Plant Sci. 14: 1219319.

Inze, D., Follin, A., Van Lijsebettens, M., Simoens, C., Genetello, C., Van Montagu, M., & Schell, J. 1984. Genetic analysis of the individual T-DNA genes of Agrobacterium tumefaciens; further evidence that two genes are involved in indole-3-acetic acid synthesis. Mol. Gen. Genet. 194: 265–274.

Joos, H., Timmerman, B., Van Montagu, M., & Schell, J. 1983. Genetic analysis of transfer and stabilization of Agrobacterium DNA in plant cells. EMBO J. 2: 2151–2160.

Killiny, N., Nehela, Y., Hajeri, S., Gowda, S., & Stelinski, L. L. 2025. Virus-induced gene silencing simultaneously exploits ‘attract and kill’ traits in plants and insects to manage Huanglongbing. Hortic Res. 12: uhae311.

Kim, H. S., Yi, H., Myung, J., Piper, K. R., & Farrand, S. K. 2008. Opine-based Agrobacterium competitiveness: Dual expression control of the agrocinopine catabolism (acc) operon by agrocinopines and phosphate levels. J. Bacteriol. 190: 3700–3711.

Kober, H., Strizhov, N., Staiger, D., Feldwisch, J., Olsson, O., Sandberg, G., Palme, K., Schell, J., & Koncz, C. 1991. T-DNA gene 5 of Agrobacterium modulates auxin response by autoregulated synthesis of a growth hormone antagonist in plants. EMBO J. 10: 3983–3991.

Koncz, C., De Greve, H., Andre, D., Deboeck, F., Van Montagu, M., & Schell, J. 1983. The opine synthase genes carried by Ti plasmids contain all signals necessary for expression in plants. EMBO J. 2: 1597–1603.

Krystel, J., Liu, H., Hartung, J., & Stover, E. 2021. Novel plantibodies show promise to protect citrus from greening disease. J. Amer. Soc. Hort. Sci. 146: 377–386.

Kwon, S. J., Bodaghi, S., Dang, T., Gadhave, K. R., Ho, T., Osman, F., Al Rwahnih, M., Tzanetakis, I. E., Simon, A. E., & Vidalakis, G. 2021. Complete nucleotide sequence, genome organization, and comparative genomic analyses of Citrus yellow-vein associated virus (CYVaV). Front. Microbiol. 12: 683130.

Liu, X., Zou, Z., Zhang, C., Liu, X., Wang, J., Xin, T., & Xia, B. 2020. Knockdown of the trehalose-6-phosphate synthase gene using RNA interference inhibits synthesis of trehalose and increases lethality rate in asian citrus psyllid, Diaphorina citri (Hemiptera: Psyllidae). Insects. 11: 605

Ma, W., Pang, Z., Huang, X., Xu, J., Pandey, S. S., Li, J., Achor, D. S., Vasconcelos, F. N. C., Hendrich, C., Huang, Y., Wang, W., Lee, D., Stanton, D., & Wang, N. 2022. Citrus Huanglongbing is a pathogen-triggered immune disease that can be mitigated with antioxidants and gibberellin. Nat. Commun. 13: 529.

Ojo, I., Ampatzidis, Y., de Oliveria Costa Neto, A., Bayabil, H. K., Schueller, J. K., & Batuman, O. 2024. The development and evolution of trunk injection mechanisms -A review. Biosyst. Eng. 240: 123–141.

Otten, L., Salomone, J. Y., Helfer, A., Schmidt, J., Hammann, P., & De Ruffray, P. 1999. Sequence and functional analysis of the left-hand part of the T-region from the nopaline-type Ti plasmid, pTiC58. Plant Mol. Biol. 41: 765–776.

Padilla, C. S., Irigoyen, S. C., Ramasamy, M., Damaj, M. B., Dominguez, M. M., Rossi, D., Bedre, R. H., Dawson, W. O., El-Mohtar, C., Irey, M. S., & Mandadi, K. K. 2025. Naturally occurring spinach defensins confer tolerance to citrus greening and potato zebra chip diseases. Plant Biotechnol. J. 23: 1876–1878.

Pradel, K. S., Ullrich, C. I., Santa Cruz, S., & Oparka, K. J. 1999. Symplastic continuity in Agrobacterium tumefaciens-induced tumours. J. Exp. Bot. 50: 183–192.

Rhodes, B. H., Stange, R. R., Zagorski, P., Hentz, M. G., Niedz, R. P., Shatters, R. G., & Pitino, M. 2023. Direct infusion device for molecule delivery in plants. J Vis Exp.

Schroder, G., Waffenschmidt, S., Weiler, E. W., & Schroder, J. 1984. The T-region of Ti plasmids codes for an enzyme synthesizing indole-3-acetic acid. Eur. J. Biochem. 138: 387–391.

Shatters, R. G., Stover, E. W., Niedz, R. P., Heck, M. L., Pitino, M., Ferrari Grando, M., & Krystal, J. (2021). Compositions and methods for modifying a plant characteristic without modifying the plant genome. WO 2021/055656 A1. March 25, 2021.

Shen, Y., Ye, T., Li, Z., Kimutai, T. H., Song, H., Dong, X., & Wan, J. 2024. Exploiting viral vectors to deliver genome editing reagents in plants. aBIOTECH. 5: 247–261.

Tipu, M. M. H., Masud, M. M., Jahan, R., Baroi, A., & Hoque, A. 2021. Identification of citrus greening based on visual symptoms: A grower’s diagnostic toolkit. Heliyon. 7: e08387.

Torti, S., Schlesier, R., Thummler, A., Bartels, D., Romer, P., Koch, B., Werner, S., Panwar, V., Kanyuka, K., Wiren, N. V., Jones, J. D. G., Hause, G., Giritch, A., & Gleba, Y. 2021. Transient reprogramming of crop plants for agronomic performance. Nat. Plants. 7: 159–171.

Ullrich, C. I., & Aloni, R. 2000. Vascularization is a general requirement for growth of plant and animal tumors. J. Exp. Bot. 51: 1951–1960.

Venkataraman, S., & Hefferon, K. 2021. Application of plant viruses in biotechnology, medicine, and human health. Viruses. 13:1697.

Wright, A. A., Shires, M. K., Beaver, C., Bishop, G., DuPont, S. T., Naranjo, R., & Harper, S. 2021. Effect of ‘Candidatus phytoplasma pruni’ infection on sweet cherry fruit. Phytopathology. 111: 2195–2202.

Ying, X., Bera, S., Liu, J., Toscano-Morales, R., Jang, C., Yang, S., Ho, J., & Simon, A. E. 2024. Umbravirus-like RNA viruses are capable of independent systemic plant infection in the absence of encoded movement proteins. PLoS Biol. 22: e3002600.

Younas, M., Zheng, Z., Atiq, M., Wei, T., & Li, Y. 2025. Insights into the Huanglongbing pandemic focusing on transmission biology, virulence factors and multifaceted management strategies. Pest Manag. Sci. 81: 6076–6094.

Zheng, D., Armstrong, C. M., Yao, W., Wu, B., Luo, W., Powell, C., Hunter, W., Luo, F., Gabriel, D., & Duan, Y. 2024. Towards the completion of Koch’s postulates for the citrus huanglongbing bacterium, Candidatus Liberibacter asiaticus. Hortic. Res. 11: uhae011.

